# AKT but not MYC promotes reactive oxygen species-mediated cell death in oxidative culture

**DOI:** 10.1101/754572

**Authors:** Dongqing Zheng, Jonathan H. Sussman, Matthew P. Jeon, Sydney T. Parrish, Alireza Delfarah, Nicholas A. Graham

## Abstract

Oncogenes can generate metabolic vulnerabilities in cancer cells. Here, we tested how AKT and MYC affect the ability of cells to shift between respiration and glycolysis. Using immortalized mammary epithelial cells, we discovered that constitutively active AKT but not MYC induced cell death in galactose culture, where cells must rely on oxidative phosphorylation for energy generation. However, the negative effects of AKT were short-lived, and AKT-expressing cells recommenced growth after ~15 days in galactose. To identify the mechanisms regulating AKT-mediated cell death, we used metabolomics and found that AKT cells dying in galactose upregulated glutathione metabolism. Next, using proteomics, we discovered that AKT-expressing cells dying in galactose upregulated nonsense-mediated mRNA decay, a marker of sensitivity to oxidative stress. We therefore measured levels of reactive oxygen species (ROS) and discovered that galactose induced ROS in cells expressing AKT but not MYC. Additionally, ROS were required for the galactose-induced death of AKT-expressing cells. We then tested whether these findings could be replicated in breast cancer cell lines with constitutively active AKT signaling. Indeed, we found that galactose induced rapid cell death in breast cancer cell lines and that ROS were required for galactose-induced cell death. Together, our results demonstrate that AKT but not MYC induces a metabolic vulnerability in cancer cells, namely the restricted flexibility to use oxidative phosphorylation.

**Implications:** The discovery that AKT but not MYC restricts the ability to utilize oxidative phosphorylation highlights that therapeutics targeting tumor metabolism must be tailored to the individual genetic profile of tumors.

## INTRODUCTION

Many cancers preferentially use glycolysis for survival and proliferation, even in the presence of oxygen, a phenomenon known as aerobic glycolysis or the Warburg effect. Increased glycolytic activity is thought to help satisfy the rapacious demands of highly proliferative cancer cells for biosynthetic precursors including lipids, proteins, and nucleic acids. However, this altered metabolism can leave tumors vulnerable to metabolic disruptions such as starvation of substrates including glucose, asparagine, glutamine, methionine, serine, and others (1–7). Therefore, understanding the interplay between oncogenes and metabolism is essential to understand how to design therapeutic strategies targeting tumor metabolism (8,9).

The altered metabolism of tumor cells has been directly linked to the same oncogenes that drive tumorigenesis. In breast cancer, both the PI3K/AKT signaling pathway and the transcription factor MYC are frequently hyperactivated (10). Deregulated PI3K/AKT signaling can result from multiple mechanisms including *PIK3CA* mutation, *PTEN* loss/mutation, or high *AKT3* expression. Hyperactivated PI3K/AKT signaling, which is prominent in the luminal subtype of breast cancer, results in increased glycolytic flux by altering the localization and activity of glycolytic enzymes including glucose transporters, hexokinase, and phosphofructokinase (11). The oncogenic transcription factor MYC is typically hyperactivated by high-level DNA amplification, especially in the basal subtype of breast cancer. Like AKT, MYC also exerts broad effects on the metabolism of tumor cells including roles in glycolysis, glutamine metabolism, nucleotide biosynthesis, and other metabolic processes (12). However, although both AKT and MYC promote aerobic glycolysis, these oncogenes can exert differential effects on metabolism in prostate cancer and pre-B cells (13,14).

The galactose culture system is a classic technique for shifting the metabolism of mammalian cells from glycolysis to aerobic respiration. In glucose-containing media, mammalian cells can utilize both glycolysis and oxidative phosphorylation (OXPHOS) to generate ATP. However, when galactose is substituted for glucose, cells must rely on OXPHOS for energy generation. Most mammalian cells exhibit a flexible metabolic state that can dynamically shift between glycolysis and OXPHOS. Because galactose culture forces oxidative metabolism, it has proved useful in the identification of inborn errors of metabolism (15,16), drugs that redirect metabolism to glycolysis (17), genes essential for OXPHOS (18), and the role of aerobic glycolysis in T cell effector functions (19). However, to our knowledge, the galactose culture system has not been used to investigate the metabolic vulnerabilities induced by oncogenes.

Here, we report that non-tumorigenic, immortalized mammary epithelial cells expressing constitutively active AKT undergo rapid cell death when switched from glucose to galactose culture. In contrast, cells expressing either RFP (i.e., negative control) or MYC readily proliferated in galactose. Cell death in AKT-expressing cells was, however, short-lived, and cells expressing AKT recommenced cell growth after ~15 days in galactose culture. To understand the differential phenotype of AKT-expressing cells, we performed metabolomic and proteomic analyses using mass spectrometry and found evidence that AKT-expressing cells in galactose were dying due to oxidative stress. We therefore used the reactive oxygen species (ROS) scavenger catalase to validate that ROS were required for galactose-induced death of AKT-expressing cells. Importantly, we also tested breast cancer cell lines with constitutively active PI3K/AKT signaling and found that these cells also exhibited cell death in galactose culture and were also rescued by catalase. Taken together, our multi-omic analysis has identified a metabolic state-dependent lethality, namely the reduced ability of cells with constitutive activation of PI3K/AKT to switch between glycolysis and respiration.

## RESULTS

### AKT but not MYC temporarily induces cell death in oxidative culture

To investigate the effect of oncogenes on the ability of human cells to survive and proliferate in oxidative culture, we first expressed either constitutively active AKT (i.e., myristoylated, or mAKT), the transcription factor MYC, or the negative control red fluorescent protein (RFP) in immortalized but non-tumorigenic MCF-10A human mammary epithelial cells. We then switched these cells from normal glucose culture to galactose culture, which forces mammalian cells to rely on oxidative phosphorylation rather the glycolysis (15). Following the switch to galactose culture, cells expressing either RFP or MYC grew ~2-fold slower than in glucose culture but did not exhibit significant cell death (Fig. 1A). In contrast, cells expressing mAKT exhibited significant cell death and declining cell numbers for three passages in galactose culture. Interestingly, mAKT-expressing cells reversed this phenotype after three passages (~15 days) and recommenced proliferation in galactose culture. However, at later passages, mAKT-expressing cells still exhibited significantly slower growth in galactose relative to glucose than did RFP- or MYC-expressing cells (Fig. 1B). Next, using liquid chromatography-mass spectrometry (LC-MS) metabolomics, we assessed the effect of galactose culture on aerobic glycolysis by measuring the intracellular concentration of lactate. We found that galactose culture significantly reduced the intracellular concentration of lactate at both short (24 h) and long (~5 passages or ~25 days) for all three cell types (Fig. 1C). Notably, mAKT-expressing cells in glucose exhibited the highest intracellular concentrations of lactate, consistent with reports that mAKT can increase aerobic glycolysis (13,20). Next, we tested whether galactose culture affected the expression of the exogenous oncogenes. After 24 h, galactose culture induced a slight decrease in expression of AKT and phospho-serine-473 AKT, but expression was restored at longer times (~5 passages) (Fig. 1D). Conversely, expression of MYC was unaffected by galactose culture. Taken together, these data demonstrate that the oncogenes mAKT and MYC exert opposite effects on the ability of non-tumorigenic MCF-10A cells to proliferate in oxidative culture conditions (i.e., galactose) at short times (< 3 passages) but that cells expressing either oncogene can proliferate in oxidative culture at longer times.

**Figure 1:**
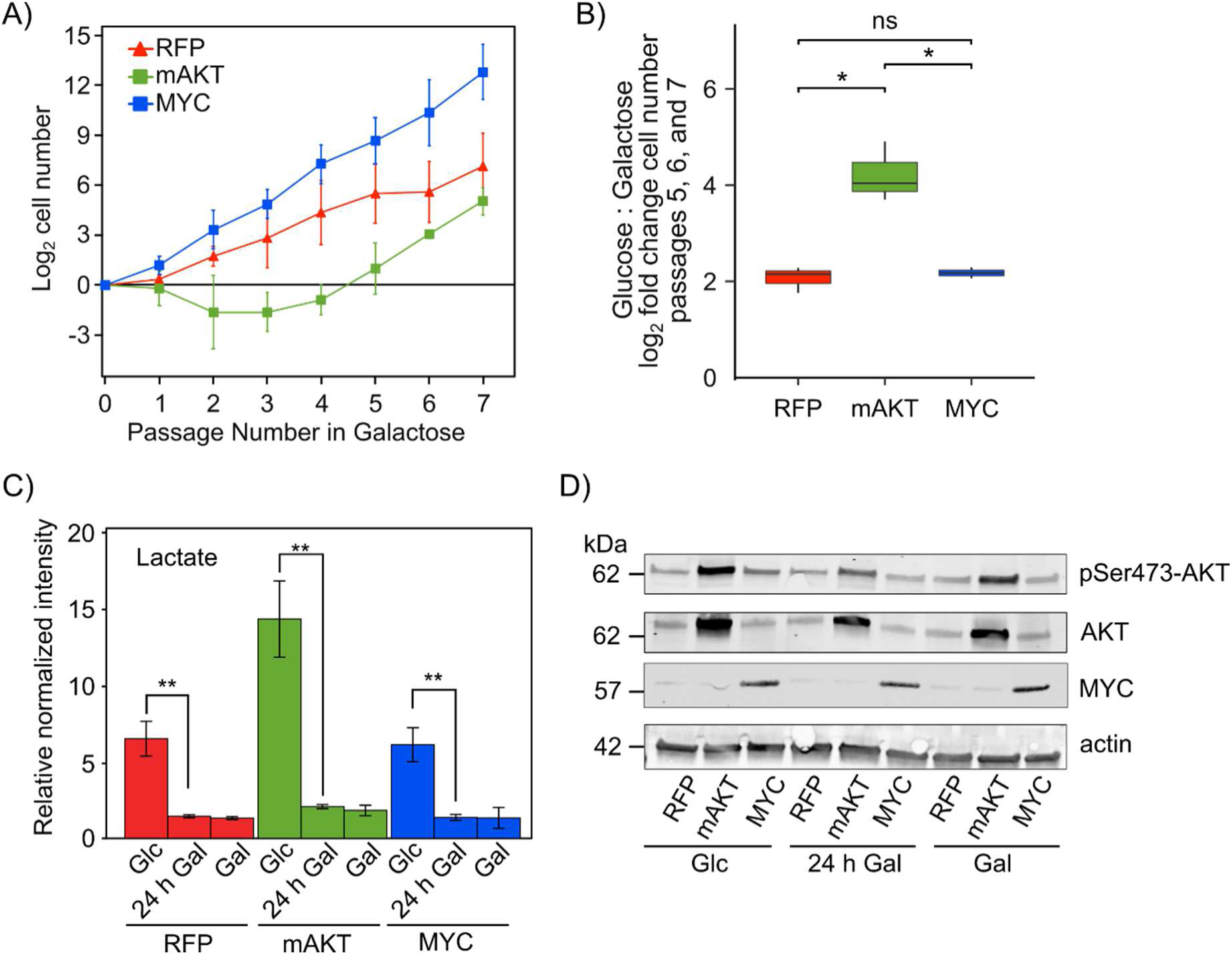
mAKT but not MYC negatively affects proliferation in galactose culture. **A)** Constitutively active (myristoylated) AKT (mAKT) but not MYC negatively affected the proliferation of MCF-10A cells grown in media containing galactose. MCF-10A cells expressing red fluorescent protein (RFP, negative control), mAKT, or MYC were switched from media containing glucose to media containing galactose and cultured for 7 passages. Galactose culture forces cells to use OXHPHOS instead of glycolysis for energy generation. For 3 passages, mAKT-expressing cells exhibited cell death and declining cell number, but afterwards, mAKT-expressing cells recommenced proliferation. The viable cell number at each passage was measured by trypan blue staining (n=2 biological replicates). **B)** mAKT-expressing cells experience a proliferative disadvantage in long-term galactose culture. A boxplot of the ratio of log_2_ fold-change for growth in glucose compared to galactose at passages 5, 6, and 7 from panel A) demonstrated that mAKT-expressing cells exhibited a growth disadvantage in galactose culture relative to RFP- or MYC-expressing cells. * denotes p-value less than 0.05 by Student’s t-test, and ns denotes not significant (n=2 biological replicates). **C)** Lactate production is suppressed in galactose culture. MCF-10A cells expressing RFP, mAKT, or MYC cells were cultured in glucose, switched from glucose to galactose for 24 h, or cultured in galactose for ~5 passages. Intracellular lactate concentrations were measured by liquid chromatography-mass spectrometry (LC-MS) metabolomics. All cell types exhibit significant reductions in intracellular lactate concentrations when cultured in galactose. mAKT-expressing cells exhibited the largest intracellular concentration of lactate in glucose culture. ** denotes p-value less than 0.01 by Student’s t-test (n=3). **D)** Expression of mAKT and MYC expression was not altered by long-term culture in galactose. MCF-10A cells expressing RFP, mAKT, or MYC were cultured in glucose, switched from glucose to galactose for 24 h, or cultured in galactose for ~5 passages, and then lysed. Protein expression was measured by Western blotting using antibodies against phospho-serine473-AKT, total AKT, or MYC. Actin was used as an equal loading control.

### mAKT-expressing cells exhibit extensive metabolic adaptation in galactose culture

To elucidate the metabolic mechanisms regulating the differential phenotypes of RFP-, mAKT-, and MYC-expressing cells in galactose culture, we profiled cells using stable isotope tracing LC-MS metabolomics (Supp. Fig. S1). Because cells in galactose culture upregulate glutamine anaplerosis (17,21), we first labeled cells with [U-^13^C]-L-glutamine for 24 h in either glucose, short-term galactose culture (24 h), or long-term galactose culture (~5 passages) and followed by LC-MS metabolomics (Supp. Fig. S2A and Supp. Table S1). Relative to RFP-expressing cells, mAKT- but not MYC-expressing cells cultured in glucose exhibited an increased percentage of M0 isotopomers for the TCA cycle intermediates citrate/isocitrate, aconitate, α-ketoglutarate, succinate, fumarate, and malate (Fig. 2A-B and Supp. Fig. S3), indicating that mAKT-expressing cells were utilizing less glutamine to fuel the TCA cycle. Upon switching to galactose culture, all cells exhibited increased percentages of fully labeled isotopomers (e.g., M6 citrate/isocitrate) indicating increased flux of glutamine-derived carbon through the TCA cycle. Additionally, in galactose culture at both short and long times, all MCF-10A cells exhibited an increased percentage of M5 isotopomer in citrate/isocitrate, indicating increased reductive carboxylation flux. Taken together, [U-^13^C]-L-glutamine stable isotope tracing indicated that all cells increased glutamine oxidation and reductive carboxylation upon switching from glucose to galactose, but that mAKT-expressing cells experienced larger changes than either RFP- or MYC-expressing cells because of their more aerobic glycolytic basal state in glucose.

**Figure 2:**
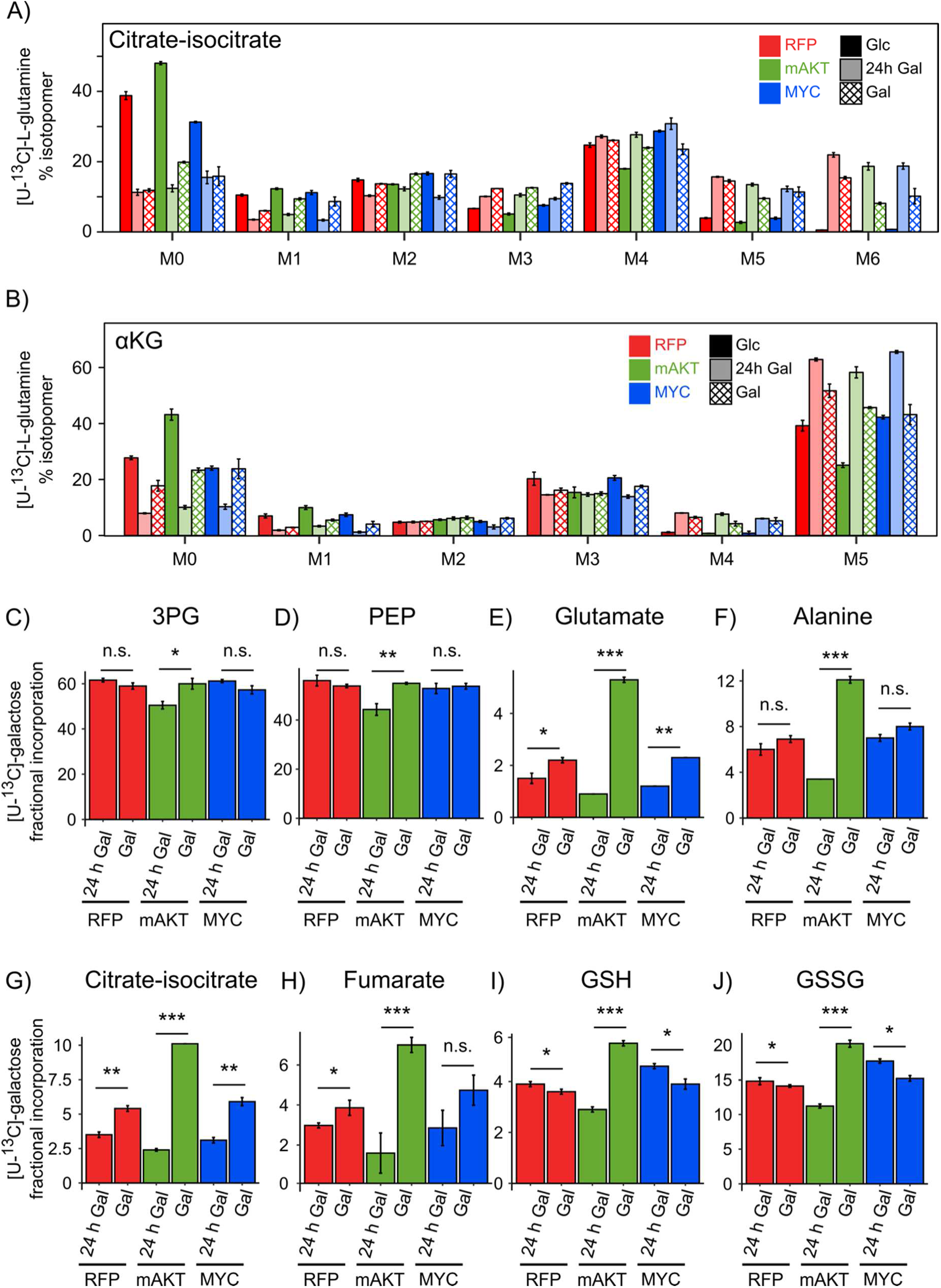
Stable isotope tracing metabolomics reveals differential usage of glutamine and galactose in mAKT-expressing cells. **A-B**) Isotopomer distributions for citrate-isocitrate and α-ketoglutarate (αKG) following [U-^13^C]-L-glutamine labeling. MCF-10A cells expressing RFP, mAKT, or MYC were cultured in glucose, switched from glucose to galactose for 24 h, or cultured in galactose for ~5 passages. Cells were labeled with [U-^13^C]-L-glutamine for 24 h and then analyzed by LC-MS metabolomics. Isotopomer abundances were normalized to total abundance to calculate the percentage of each isotopomer. mAKT-expressing cells in glucose exhibited a larger percentage of M0 isotopomer than either RFP- or MYC-expressing cells. All cell types exhibited more reductive carboxylation (e.g., M5 citrate-isocitrate) and more glutamine anaplerosis (e.g., M6 citrate-isocitrate) in galactose culture. **C-J**) Fractional contribution for MCF-10A cells labeled with [U-^13^C]-galactose for selected metabolites from glycolysis (C, D), amino acid metabolism (E, F), the TCA cycle (G, H), and glutathione metabolism (I, J). MCF-10A cells expressing RFP, mAKT, or MYC were switched from glucose to galactose for 24 h or cultured in galactose for ~5 passages. Cells were labeled with [U-^13^C]-galactose for 24 h and then analyzed by LC-MS metabolomics. 3PG denotes 3-phosphoglycerate, PEP denotes phosphoenolpyruvate, GSH denotes reduced glutathione, and GSSG denotes oxidized glutathione. mAKT-expressing cells exhibited larger changes in ^13^C fractional incorporation than either RFP- or MYC-expressing cells. * denotes p-value less than 0.05, ** denotes p-value less than 0.01, *** denotes p-value less than 0.001 by Student’s t-test, and n.s. denotes not significant (n=3).

Next, we tested how MCF-10A cells expressing oncogenes adapt to galactose culture by labeling cells with [U-^13^C]-galactose in short-term (24 h) and long-term galactose culture (~5 passages) followed by LC-MS metabolomics (Supp. Fig. S2B and Supp. Table S2). Examining the fractional incorporation of ^13^C, we found that mAKT-expressing cells exhibited a small but significant reduction in ^13^C fractional incorporation compared to RFP- and MYC-expressing cells in the glycolytic intermediates 3-phosphoglycerate (3PG) and phosphoenolpyruvate (PEP), indicating slower flux from galactose into glycolysis (Fig. 2C-D). At longer times, however, mAKT cells reversed this difference and exhibited ^13^C fractional incorporation to levels similar to RFP- and MYC-expressing cells, indicative of increased flux from galactose into glycolysis. In addition, we found that mAKT-expressing cells significantly increased the ^13^C fractional incorporation from galactose into the amino acids glutamate and alanine (Fig. 2E-F), TCA cycle intermediates including citrate/isocitrate and fumarate (Fig. 2G-H), and the redox buffering molecules reduced (GSH) and oxidized glutathione (GSSG) (Fig. 2I-J). In contrast, RFP- and MYC-expressing cells exhibited smaller increases in ^13^C fractional incorporation in glutamate, alanine, citrate/isocitrate, and fumarate, and decreased ^13^C fractional incorporation into GSH and GSSG. Overall, the changes in galactose-derived ^13^C fractional incorporation for mAKT-expressing cells were much greater than for RFP- or MYC-expressing cells. Taken together, this demonstrates that AKT-expressing cells exhibited significantly larger adaptations to galactose culture than do RFP- or MYC-expressing cells, consistent with the observation that AKT-expressing cells initially die in galactose culture before recommencing proliferation.

### Short-term galactose culture increases glutathione metabolism in mAKT-expressing cells

To further understand the metabolic profile that occurs in MCF-10A cells expressing oncogenes when switched from glucose to galactose, we next analyzed metabolite pool sizes (Supp. Table S3). Unsupervised hierarchical clustering of metabolite pool sizes segregated samples based on the culture condition (glucose, short-term galactose, and long-term galactose culture) rather than by oncogene (RFP, mAKT, and MYC) (Fig. 3A). No obvious grouping of functionally related metabolites was apparent from the hierarchical clustering of metabolites. Therefore, to identify metabolic pathways affected by galactose culture in each cell type, we employed Metabolite Set Enrichment Analysis (MSEA) (13,22,23). First, we analyzed the relative enrichment of all KEGG pathways comparing short-term galactose to glucose culture. To identify differentially enriched pathways, we plotted the MSEA results on a volcano-style plot (Fig. 3B). This analysis revealed that glutathione metabolism was significantly enriched in short-term galactose mAKT- but not RFP- or MYC-expressing cells (Fig. 3C and Supp. Fig. S4A,C). Next, we conducted a similar analysis comparing metabolite pool sizes in short-term and long-term galactose cultured cells. Again, we found that glutathione metabolism was significantly enriched in short-term galactose mAKT- but not RFP- or MYC-expressing cells (Fig. 3D-E and Supp. Fig. S4B,D). Taken together, this metabolite set enrichment analysis suggests that glutathione metabolism is specifically upregulated in mAKT-expressing cells when initially switched to galactose culture.

**Figure 3:**
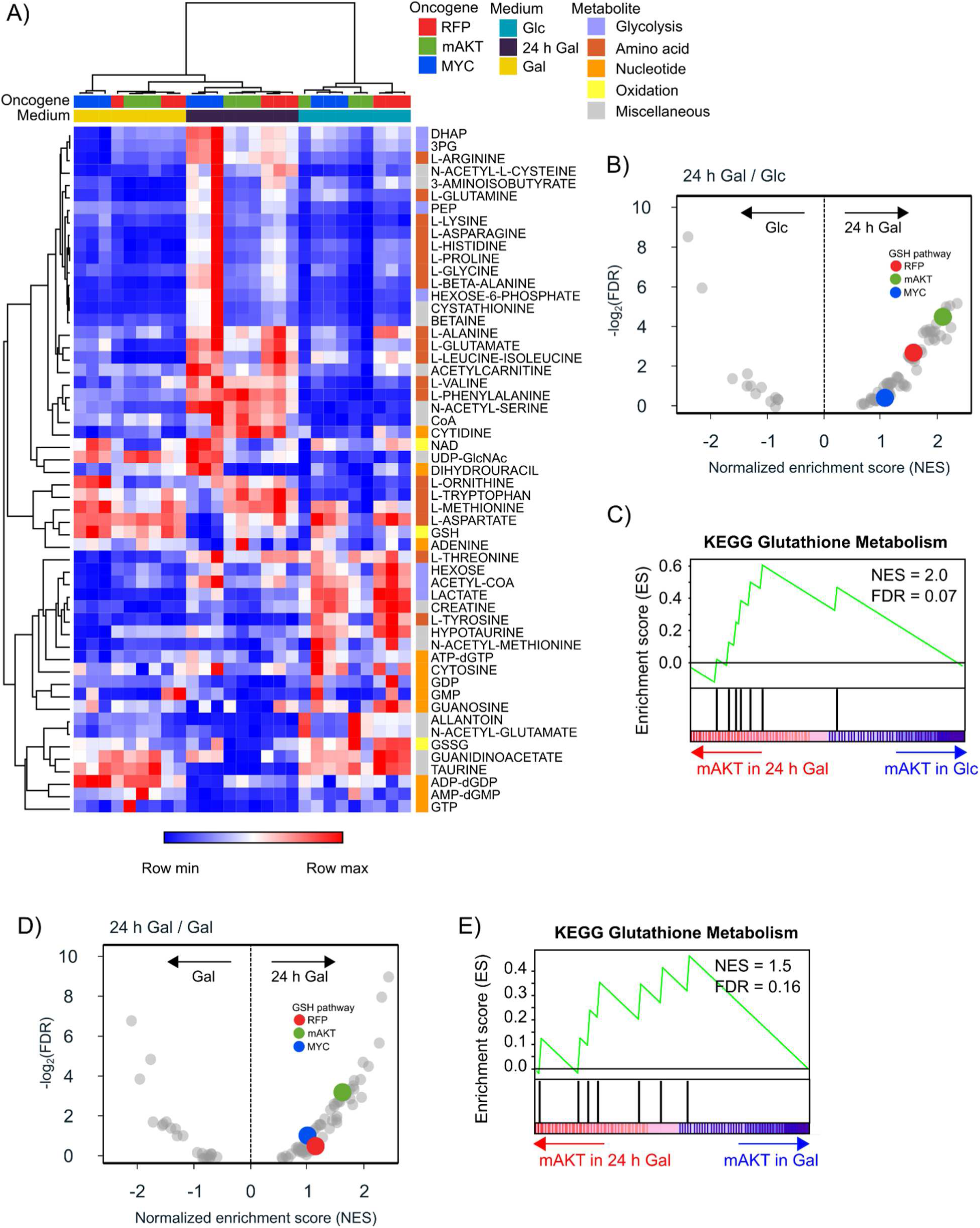
Glutathione metabolism is enriched in mAKT- but not RFP- or MYC-expressing MCF-10A cells in short-term galactose culture. **A**) Hierarchical clustering of metabolite pool sizes separated samples by media type but not oncogene. MCF-10A cells expressing RFP, mAKT, or MYC were cultured in glucose (Glc), switched from glucose to galactose for 24 h (24 h Gal), or cultured in galactose for ~5 passages (Gal), and metabolite pool sizes were measured by LC-MS metabolomics. Metabolite pool sizes were filtered for ANOVA p-value < 0.5, and clustered using one minus the Pearson correlation and average linkage. Samples clustered by media type (e.g., Glc) rather than by oncogenes (e.g., RFP). Metabolites are colored at right of the heatmap according to their metabolic pathways. Red and blue denote higher and lower abundance, respectively, from samples run in biological triplicate. **B, C**) Metabolite set enrichment analysis of metabolite pool sizes from short-term galactose culture (24 h Gal) relative to glucose culture (Glc). All KEGG metabolic pathways were analyzed. B) Volcano plot of -log_2_(false discovery rate, FDR) vs. the normalized enrichment score (NES). The glutathione metabolism pathway (GSH pathway) for cells expressing RFP, mAKT, and MYC is highlighted. All other pathways are shown in light gray. C) Mountain plot of KEGG Glutathione Metabolism demonstrating that glutathione metabolism was significantly enriched in mAKT-expressing cells following 24 h of culture in galactose. The green line denotes the enrichment score, and the black tick marks denote metabolites that belong to glutathione metabolism. **D, E**) Metabolite set enrichment analysis of metabolite pool sizes from short-term galactose culture (24 h Gal) relative to long-term galactose culture (~5 passages, Gal). All KEGG metabolic pathways were analyzed. D) Volcano plot of -log_2_(FDR) vs. the normalized enrichment score (NES). The glutathione metabolism pathway (GSH pathway) for cells expressing RFP, mAKT, and MYC is highlighted. All other pathways are shown in light gray. C) Mountain plot of KEGG Glutathione Metabolism demonstrating that glutathione metabolism was significantly enriched in mAKT-expressing cells following 24 h of culture in galactose. The green line denotes the enrichment score, and the black tick marks denote metabolites that belong to glutathione metabolism.

### Proteomic analysis reveals enriched nonsense-mediated mRNA decay (NMD) in mAKT-expressing cells in short-term galactose culture

Next, to further characterize the mechanisms underlying the differential phenotype of mAKT-expressing cells in galactose culture, we performed label-free quantitative LC-MS proteomics. We analyzed two independent biological experiments of MCF-10A cells expressing either RFP, mAKT, or MYC cultured in glucose, short-term galactose, and long-term galactose culture. Across both experiments, we identified and quantified 2,460 proteins, 1,356 of which were quantified in both biological replicates (Fig. 4A and Supp. Table S4). To identify global differences between samples, we conducted principal component analysis (PCA) using the proteins identified in both biological replicates. PCA revealed a clear separation across samples and consistent trends across the two experiments. Notably, in both replicates, short-term galactose culture induced a positive shift on PC1 (48.4% of variation for Experiment 1) for all cell types (Fig. 4B and Supp. Fig. S5A). Long-term galactose culture, in contrast, generally exhibited a negative shift on PC1 relative to short-term galactose culture. Notably, the PC1 shift for mAKT-expressing cells was significantly larger than for either RFP- or MYC-expressing cells. To understand the proteins driving separation on PC1, we analyzed the PC1 loadings vector using 1D annotation enrichment using the Reactome Pathway Database (24,25). Among the enriched pathways, we found that two nonsense-mediated mRNA decay (NMD) pathways were enriched in positive PC1, in addition to several other mRNA-related pathways (Fig. 4C and Supp. Fig. S5B). To better understand how these pathways were regulated by galactose culture, we examined members of Nonsense-mediated mRNA decay (NMD) enhanced by the Exon Junction Complex (EJC), a surveillance pathway that eliminates mRNAs containing premature translation stop codons (26). Visualization of these proteins on a heatmap revealed that their basal expression was low in mAKT-expressing cells cultured in glucose and that short-term galactose induced higher expression (Fig. 4D and Supp. Fig. S5C). In long-term galactose culture, the levels of NMD proteins were again reduced to low expression in mAKT-expressing cells. Taken together, this proteomic analysis suggests that mAKT-expressing cells in short-term galactose culture dramatically upregulated NMD.

**Figure 4:**
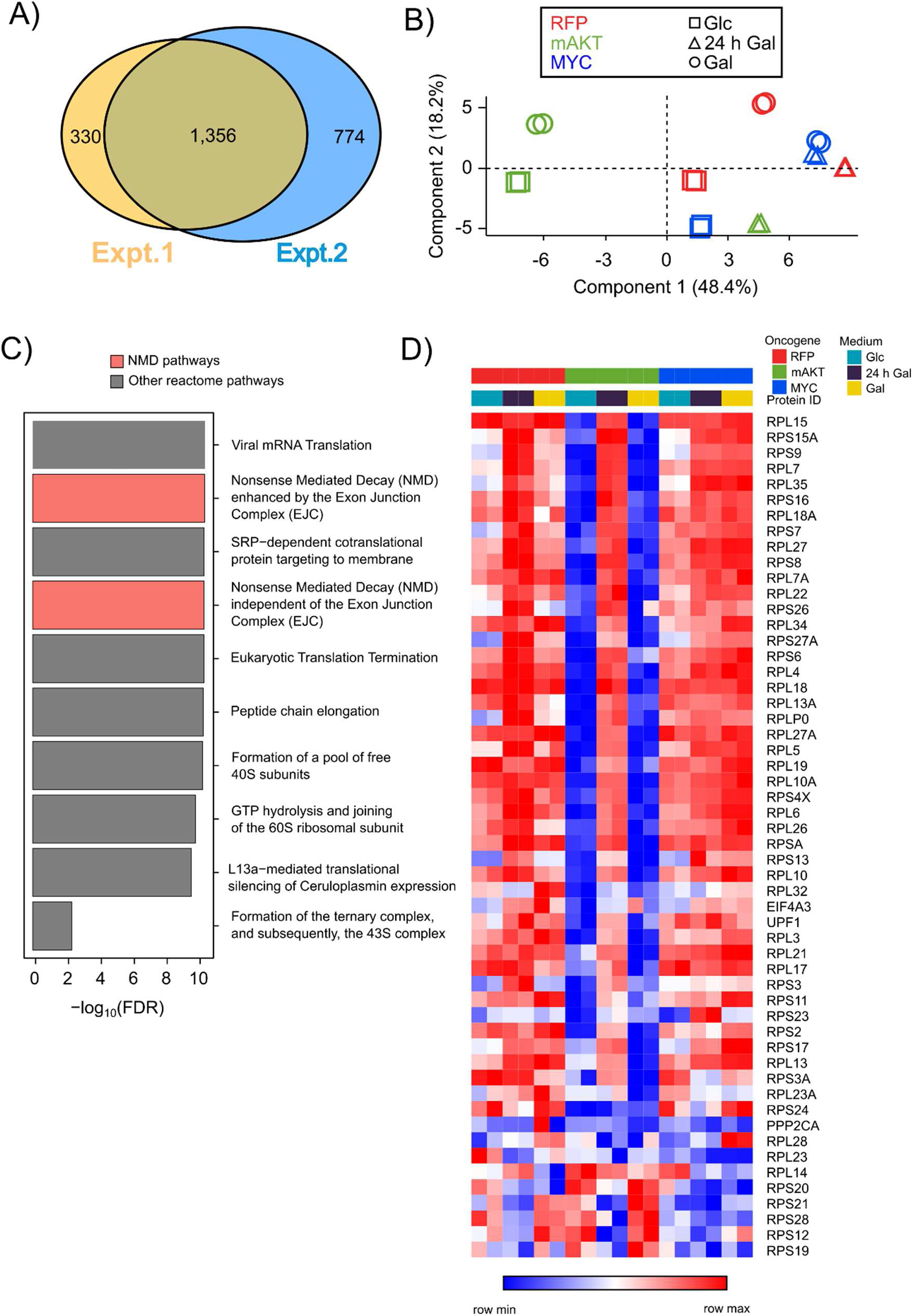
Proteomic profiling reveals that mAKT-expressing cells exhibit upregulation of nonsense-mediated mRNA decay (NMD) proteins in short-term galactose culture. **A)** Overlap in quantified proteins by LC-MS proteomics from two independent biological replicates. MCF-10A cells expressing RFP, mAKT, or MYC were cultured in glucose (Glc), switched from glucose to galactose for 24 h (24 h Gal), or cultured in galactose for ~5 passages (Gal), and protein expression was measured by label-free LC-MS proteomics. Experiments 1 and 2 identified and quantified 1,686 and 2,130 proteins, respectively. 1,356 proteins were quantified in both experiments in technical duplicate. **B)** Principal component analysis score plots (PC1 vs. PC2) of proteomic data from Experiment 1 segregated samples by oncogene and media type. Only proteins identified in both experiments were used. Color denotes oncogene, and shape denotes media type. Each sample was analyzed in technical duplicate. Short-term galactose culture (24 h Gal) induced a positive shift on PC1 (48.4% of variation) for all cell types relative to glucose culture (Glc). Long-term galactose culture (Gal), in contrast, exhibited a negative shift on PC1 relative to short-term galactose culture for mAKT- and RFP-expressing cells. The PC1 shift for mAKT-expressing cells was significantly larger than for either RFP- or MYC-expressing cells. Similar trends were seen in Experiment 2 (Supp. Fig. S5A). **C)** Enrichment analysis identified Reactome pathways enriched in the PC1 loadings vector from Experiment 1. Nonsense Mediated Decay (NMD) enhanced by the Exon Junction Complex (EJC) and Nonsense Mediated Decay (NMD) independent of the Exon Junction Complex (EJC) are highlighted. Similar trends were seen in Experiment 2 (Supp. Fig. S5B). **D)** mAKT-expressing cells significantly upregulated NMD proteins in short-term galactose culture. A heatmap of protein expression from Experiment 1 for the Nonsense Mediated Decay (NMD) enhanced by the Exon Junction Complex (EJC) pathway (Reactome R-HSA-975957) demonstrated that mAKT-expressing cells dramatically upregulated NMD protein expression in short-term (24 h) galactose culture. Similar trends were seen in Experiment 2 (Supp. Fig. S5C).

### mAKT but not RFP- or MYC-expressing cells exhibited oxidative stress in short-term galactose culture

In mammalian cells, activation of NMD can sensitize cells to oxidative stress (27). Because our metabolomic analysis (Figs. 3,4) demonstrated that mAKT-expressing cells in short-term galactose culture upregulated glutathione metabolism, we hypothesized that AKT-expressing cells were dying from increased oxidative stress following the switch from glucose to galactose culture. We thus measured the levels of reactive oxygen species (ROS) using the fluorescent ROS probe DCF-DA. MCF-10A cells expressing either RFP, mAKT, or MYC were switched from glucose to galactose culture for 3 h and ROS levels were measured by flow cytometry. In RFP- and MYC-expressing cells, the switch to galactose culture did not significantly alter ROS levels (Fig. 5). In mAKT-expressing cells, however, 3 h of galactose culture increased the levels of ROS approximately ten-fold. Thus, constitutively active AKT signaling but not MYC induced elevated ROS in galactose-cultured MCF-10A cells.

**Figure 5:**
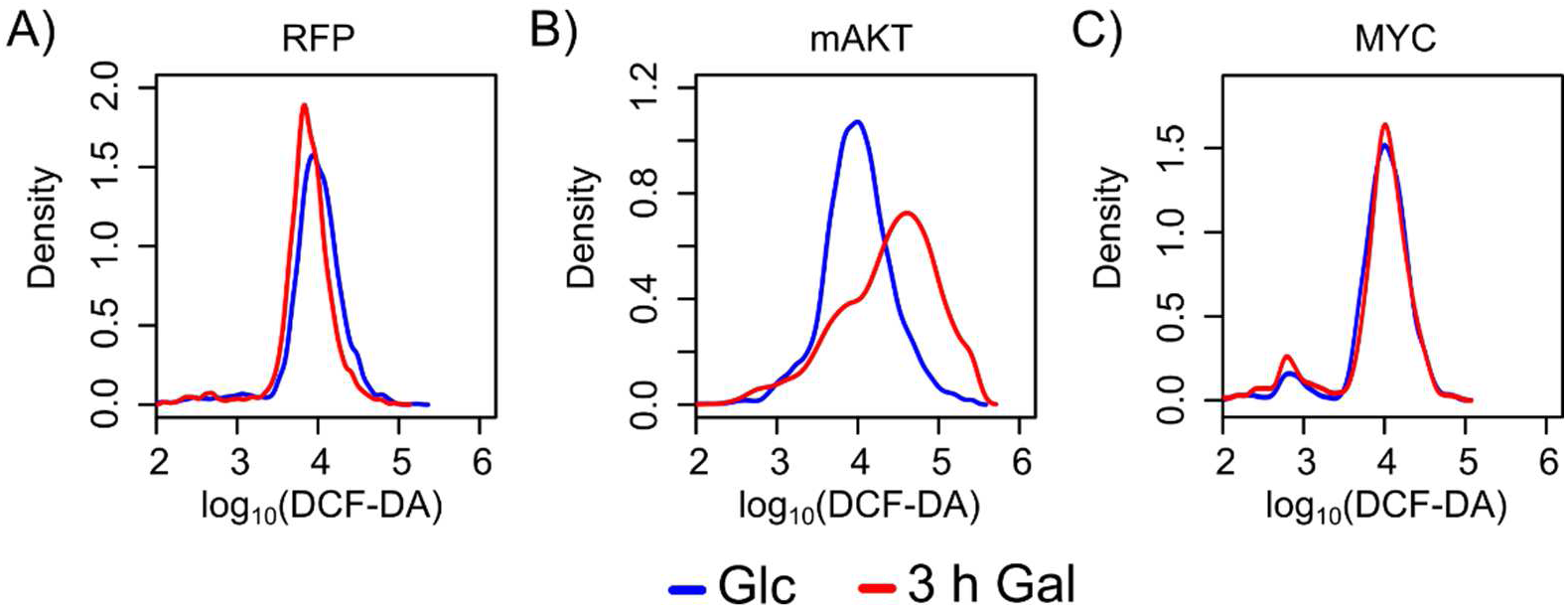
mAKT- but not RFP- or MYC-expressing cells exhibited increased ROS upon switching from glucose to galactose. **A-C**) MCF-10A cells expressing RFP, mAKT, or MYC were cultured in glucose or switched from glucose to galactose for 3 h (3 h Gal). ROS levels were measured by flow cytometry using the fluorescent ROS probe DCF-DA. Only mAKT-expressing cells exhibited increased levels of ROS following galactose culture.

### Galactose culture-induced cell death can be rescued by ROS scavengers

Having identified that mAKT-expressing cells experience oxidative stress in short-term galactose culture, we next tested whether ROS were functionally involved in the cell death caused by galactose culture. We cultured MCF-10A cells expressing either RFP, mAKT, or MYC in media without glucose, media with galactose, or media with galactose plus the ROS scavenger catalase. For MCF-10A cells expressing either RFP or mAKT, glucose starvation resulted in significant cell death (Fig. 6A and Supporting Fig. S6A). Interestingly, cells expressing MYC were protected from glucose starvation-induced cell death, mirroring observations in normal human fibroblasts and glioma cells (6,7). In galactose culture, however, only MCF-10A cells expressing mAKT exhibited cell death. In these mAKT-expressing cells, supplementation with the ROS scavenger catalase rescued cells from galactose culture-induced cell death. Thus, ROS induced by galactose culture are required for galactose culture-induced cell death.

**Figure 6:**
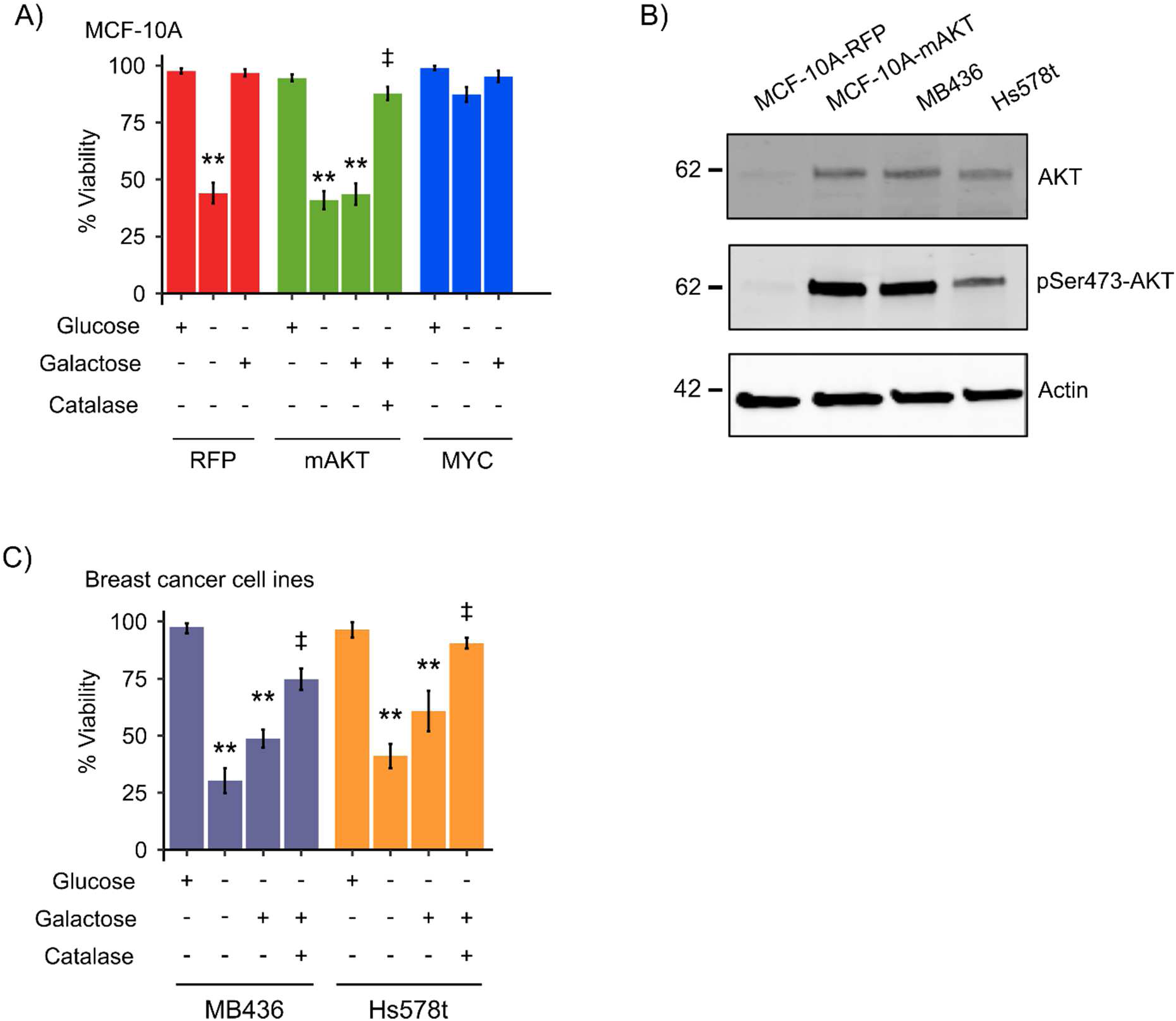
ROS are required for galactose-induced cell death. **A)** The ROS scavenger catalase rescued mAKT-expressing cells from galactose-induced cell death. MCF-10A cells expressing RFP, mAKT, or MYC were cultured in glucose, without glucose, or in galactose with or without 200 U/ml of the ROS scavenger catalase for 32 h. Cell viability was measured by trypan blue staining. ** denotes Student’s t-test p-value < 0.01 compared to glucose culture, and ‡ denotes Student’s t-test p-value < 0.01 compared to galactose culture without catalase (n=2-4 biological replicates). **B)** M436 and Hs578t breast cancer cells exhibit constitutively active PI3K/AKT signaling. MCF-10A cells expressing either RFP or mAKT and M436 and Hs578t breast cancer cell lines were serum starved for 16 h and then lysed. Western blotting with antibodies against total AKT and phosphoSer473-AKT demonstrated that MCF-10A expressing mAKT and the breast cancer cell lines exhibited constitutively active AKT. Actin was used as an equal loading control. **C)** The ROS scavenger catalase rescued MB436 and Hs578t breast cancer cells from galactose-induced cell death. MB436 and Hs578t cells were cultured in glucose, without glucose, or in galactose with or without 200 U/ml of the ROS scavenger catalase for 32 h. Cell viability was measured by trypan blue staining for 24 h, and cell viability was measured by trypan blue staining. ** denotes Student’s t-test p-value < 0.01 compared to glucose culture, and ‡ denotes Student’s t-test p-value < 0.01 compared to galactose culture without catalase (n=3-4 biological replicates).

Next, we sought to determine if our results in MCF-10A expressing mAKT were also reflected in breast cancer cell lines. We chose two cell lines reported to exhibit constitutively active AKT signaling, MDA-MB-436 (MB436) and Hs578t (28,29). Western blotting confirmed that both cell lines exhibited AKT activation even after serum starvation (Fig. 6B). Next, we tested the effect of glucose starvation and galactose culture on these cell lines and found that both cell lines exhibited significant cell death upon glucose starvation and short-term galactose culture (Fig. 6C and Supp. Fig. S6B). In addition, for both cell lines, the ROS scavenger catalase rescued cells from galactose culture-induced cell death. Taken together, these results show that breast cancer cell lines with constitutively active AKT signaling experience ROS-mediated cell death when cultured in galactose medium, similar to our MCF-10A-mAKT cells.

## DISCUSSION

The altered metabolism of tumors has long been proposed as a therapeutic target. Defining the metabolic vulnerabilities induced by specific oncogenes is crucial for the design and stratification of therapeutics targeting tumor metabolism (8,9). Here, using the galactose culture system, which forces mammalian cells to rely on OXPHOS instead of glycolysis for energy generation (15–19), we have uncovered a metabolic state-dependent lethality, namely the restricted ability of cells with constitutively active PI3K/AKT signaling to switch between glycolysis and OXPHOS. Through multi-omic analysis, we identified that the galactose-induced cell death exhibited by mAKT-expressing cells was accompanied by increased glutathione metabolism (Fig. 3) and increased expression of NMD proteins (Fig. 4), a hallmark of sensitivity to oxidative stress. Based on these results, we found that galactose-induced cell death required ROS, both in MCF-10A cells expressing mAKT and in breast cancer cell lines with activated PI3K/AKT signaling (Fig. 5-6). Taken together, these results reveal a novel metabolic state vulnerability induced by PI3K/AKT signaling.

Our findings in MCF-10A, a non-tumorigenic but immortalized breast cancer cell line, confirm reports that constitutive activation of PI3K/AKT signaling forces cells to rely on aerobic glycolysis (14,20). Stable isotope tracing with [U-^13^C]-L-glutamine confirmed that these cells exhibit less glutamine anaplerosis than either RFP- or MYC-expressing cells when cultured in glucose (Fig. 2A, B). Interestingly, MCF-10A cells expressing the negative control RFP were sensitive to glucose deprivation, and mAKT expression did not further sensitize cells to glucose starvation (Fig. 6A). However, RFP-expressing cells were able to switch from glycolysis to aerobic respiration, as evidenced by their survival and growth in galactose media (Figs. 1, 6). mAKT-expressing cells, in contrast, were initially unable to metabolize galactose, leading to ROS-mediated cell death even though PI3K/AKT activation can increase resistance to oxidative stress by upregulating glutathione biosynthesis (30). In addition, our observation that MYC did not sensitize MCF-10A cells to galactose culture supports previous reports that AKT and MYC differentially alter metabolism, including the creation of metabolic vulnerabilities in glycolysis and mitochondrial bioenergetics, respectively (13,14). Similarly, our data confirm that MYC activation protects against glucose deprivation (6,7), though MYC expression in MCF-10A cells did not exhibit substantial increases in oxidative metabolism relative to RFP-expressing cells as has been seen in other cell types (31). Together, the differential sensitivity of AKT- and MYC-expressing cells suggests that these oncogenes alter metabolism through distinct mechanisms even though both can upregulate aerobic glycolysis (11,12,32).

Although mAKT-expressing cells initially die in galactose culture, these cells recommence proliferation after ~15 days. At present, it is not known if this represents metabolic remodeling to process galactose (i.e., acquired resistance) or the emergence of a pre-existing sub-population of galactose-metabolizing cells (i.e., clonal selection). It is notable, however, that the AKT-expressing cells proliferating in galactose exhibit slower growth relative to glucose than either RFP- or MYC-expressing cells (Fig. 1B). Thus, even the “galactose-resistant” mAKT-expressing cells exhibit a disadvantage in oxidative culture, which may explain why proliferating mAKT cells exhibit a substantially different galactose metabolism than RFP- or MYC-expressing cells (Fig. 2E-F). Notably, because long-term culture in galactose did not alter AKT expression or phosphorylation, the changes that enable proliferation of mAKT-expressing cells in galactose must occur downstream of AKT itself (Fig. 1D). These changes could include increased mitochondrial content and aerobic metabolism, as has been observed when human myotubes, which have low mitochondrial oxidative potential, are cultured in galactose (33,34). Regardless, the fact that mAKT cells reliably and rapidly demonstrates resistance to galactose suggests that eradication of tumors with constitutive PI3K/AKT signaling will require the therapeutic targeting of another, compensatory pathway to prevent re-emergence. Notably, at this time, we have been unable to generate galactose-resistant clones of either MB436 or Hs578t, suggesting that these breast cancer cells harbor oncogenic lesions in addition to PI3K/AKT signaling that more permanently restrict the flexibility to utilize OXPHOS.

Another possible mechanism by which mAKT-expressing cells adapt to galactose culture might be through suppression of NMD (Fig. 4). In general, oxidative stress suppresses NMD in order to enhance the ability of cells to survive ROS toxicity (26,27). However, when NMD is activated, cells become more sensitive to oxidative stress (27). Thus, the galactose-induced cell death of mAKT-expressing may be due to increased ROS levels coinciding with increased sensitivity to oxidative stress due to NMD upregulation. In fact, there may exist a direct link between AKT signaling and NMD upregulation, as insulin signaling in HeLa cells upregulates NMD through increased binding of UPF1, the master regulator of NMD, to mRNA transcripts (35). In our MCF-10A-AKT and breast cancer cell lines, the mechanisms by which AKT signaling in galactose induces the upregulation of NMD proteins are currently under investigation.

In summary, the deregulation of PI3K/AKT and MYC in breast cancer motivates further research into how these oncogenes generate oncogene-dependent metabolic vulnerabilities. Like AKT and MYC, many other frequently altered oncogenes including BRAF (36), ERBB2 (37), KRAS (38,39), and VHL (40) alter the balance between glycolysis and OXPHOS. However, it remains to be tested whether these oncogenes will affect the flexibility of cells to shift between respiration and OXPHOS as does AKT. Regardless of the oncogene, it will be crucial to understand how oncogenic events define the response to fluctuations in nutrient availability, oxygen tension, and pH within the tumor microenvironment (41). In addition, given the flexibility and adaptability of cancer metabolism, inhibition of a single molecular target (e.g., glycolysis in tumors with hyperactivated AKT signaling) may not prove sufficient for tumor eradication. As such, combinational therapies that generate synthetic lethality in tumors also need to be investigated in the context of oncogene dependence (42–44). Taken together, our findings highlight that the importance of oncogene-dependent metabolic vulnerabilities in cancer cells and suggest that therapies targeting tumor metabolism will need to be appropriately paired with tumor genetic profiles.

## MATERIALS AND METHODS

### Cell culture

MCF-10A human mammary epithelial cells were obtained from American Type Culture Collection (ATCC, obtained in November 2016). Cells were cultured in DMEM/Ham’s F-12 supplemented with 5% horse serum, 100 ng/ml cholera toxin, 20 ng/ml epidermal growth factor, 10 µg/ml insulin, 500 ng/ml hydrocortisone, 1% penicillin/streptomycin/amphotericin B, and either 25 mM glucose or galactose. MDA-MB-436 (MB436) and Hs578t cells were a gift from Dr. Michael Press (USC Department of Pathology, obtained in January 2019). MB436 and Hs578t cells were cultured in DMEM supplemented with 10% FBS and 1% penicillin/streptomycin/amphotericin B. Dialyzed FBS was used for all experiments where MB436 and Hs578t cells were cultured with galactose. All cells were grown in a 5% CO_2_, 37°C, and humidified incubator and were used within 30 passages of thawing. Cell counting and viability were assessed using trypan blue staining with a TC20 automated cell counter (BioRad).

### Retroviral infection

RFP, myristoylated AKT, and MYC were cloned into the pDS-FB-neo retroviral vector and verified by Sanger sequencing. Retrovirus was prepared in 293T cells by co-transfection with viral packaging plasmids. Following infection, MCF-10A cells were selected with 1 mg/ml G418. Following selection, cells were maintained in media with 500 µg/ml G418.

### Reactive oxygen species measurement

MCF-10A were treated in respective media for 3 h. Hydrogen peroxide (1 mM) was used as a positive control. The cells were incubated with 5 µM of DCF-DA (Biotium #10058) for 30 min at 37°C MCF-10A prior to trypsinization. The cells were washed with PBS twice, lifted with TrypLE, and resuspended in PBS for an approximate final concentration of 1 million cells/ml. The ROS probed samples were analyzed on a Miltenyi Biotec MACSQuant flow cytometer with FITC channel (488 nm excitation/520 nm emission) to measure fluorescence, and data were processed and analyzed with flowCore (1.48.1) R package.

### Western blotting

Cells were lysed in modified RIPA buffer (50 mM Tris–HCl (pH 7.5), 150 mM NaCl, 50 mM β-glycerophosphate, 0.5 mM NP-40, 0.25% sodium deoxycholate, 10 mM sodium pyrophosphate, 30 mM sodium fluoride, 2 mM EDTA, 1 mM activated sodium vanadate, 20 µg/ml aprotinin, 10 µg/ml leupeptin, 1 mM DTT, and 1 mM phenylmethylsulfonyl fluoride). Whole-cell lysates were resolved by SDS–PAGE on 4–15% gradient gels and blotted onto nitrocellulose membranes (Bio-Rad). Membranes were blocked for 1 h and then incubated with primary overnight and secondary antibodies for 2 h. Blots were imaged using the Odyssey Infrared Imaging System (Li-Cor). Primary antibodies used for Western blot analysis were: anti-AKT (Cell Signaling Technology 9272), anti-phospho-Ser473-AKT (Santa Cruz Biotechnology sc-7985-R), anti-c-MYC (Cell Signaling Technology 9402), and anti-β-actin (Proteintech 66009-1-lg).

### LC-MS Metabolomics

MCF-10A cells were plated on 6-well plates at the density of 7,333 cells/cm^2^. After 24 h, media was removed, cells were washed twice with 2 mL of PBS, and 1 mL of media was added to cells. Media contained either [U-^13^C]-L-glutamine or [U-^13^C]-galactose (Cambridge Isotope Laboratories). After 24 h, the culture plates were cooled on ice, media was aspirated, and the cells were washed with 1 mL of cold ammonium acetate. Upon aspirating the ammonium acetate, metabolites were extracted with 1 mL of −80°C methanol. The methanol cell suspension was scraped and transferred to Eppendorf tubes, and the cell suspension was centrifuged at 4°C. The supernatants was transferred to a new Eppendorf tubes, and the pellet was re-extracted with another 350 µL of −80°C methanol. The second methanol extraction was spun down, and the supernatant was pooled with the first extraction. Metabolites were speed-vac dried, resuspended in LC-MS grade water, and sent for LC-MS analysis.

Samples were randomized and analyzed on a Q Exactive Plus hybrid quadrupole-Orbitrap mass spectrometer coupled to an UltiMate 3000 UHPLC system (Thermo Scientific). The mass spectrometer was run in polarity switching mode (+3.00 kV/-2.25 kV) with an m/z window ranging from 65 to 975. Mobile phase A was 5 mM NH_4_AcO, pH 9.9, and mobile phase B was acetonitrile. Metabolites were separated on a Luna 3 µm NH2 100 Å (150 × 2.0 mm) column (Phenomenex). The flowrate was 300 µL/min, and the gradient was from 15% A to 95% A in 18 min, followed by an isocratic step for 9 min and re-equilibration for 7 min. All samples were run in biological triplicate. Metabolites were detected and quantified as area under the curve based on retention time and accurate mass (≤ 8 ppm) using the TraceFinder 3.3 (Thermo Scientific) software. Raw data were corrected for naturally occurring ^13^C abundance (45). Intracellular data was normalized to the cell number at the time of extraction.

### LC-MS Proteomics

MCF-10A cells dishes were placed on ice and washed with PBS. Cells were then scraped and pelleted by centrifugation. The cell pellets were lysed by probe sonication in 8 M urea (pH 7.5), 50 mM Tris, 1 mM activated sodium vanadate, 2.5 mM sodium pyrophosphate, 1 mM β-glycerophosphate, and 100 mM sodium phosphate. The above procedures were performed in 0-4°C. Insoluble cell debris were filtered by 0.22 um syringe filter. Protein concentration was measured by BCA assay (Pierce, PI23227). Lysates were reduced with 5 mM DTT, alkylated with 25 mM iodoacetamide, quenched with 10 mM DTT, and acidified to pH 2 with 5% trifluoracetic acid. Proteins were then digested to peptides using a 1:100 trypsin to lysate ratio by weight. Tryptic peptides were desalted by reverse phase C18 StageTips and eluted with 30% acetonitrile. The eluents were vacuumed dried, and 250 ng/injection was submitted to LC-MS. We performed two independent biological replicates, and each experiment were subjected to two technical LC-MS injections.

The samples were randomized and injected into an Easy 1200 nanoLC ultra high-performance liquid chromatography coupled with a Q Exactive quadruple orbitrap mass spectrometry (Thermo Fisher). Peptides were separated by a reverse-phase analytical column (PepMap RSLC C18, 2 µm, 100Å, 75 µm X 25 cm). Flow rate was set to 300 nL/min at a gradient from 3% buffer B (0.1% formic acid, 80% acetonitrile) to 38% B in 110 min, followed by a 10-minute washing step to 85% B. The maximum pressure was set to 1,180 bar and column temperature was maintained at 50°C. All samples were run in technical duplicate. Peptides separated by the column were ionized at 2.4 kV in the positive ion mode. MS1 survey scans were acquired at the resolution of 70k from 350 to 1800 m/z, with maximum injection time of 100 ms and AGC target of 1e6. MS/MS fragmentation of the 14 most abundant ions were analyzed at a resolution of 17.5k, AGC target 5e4, maximum injection time 65 ms, and normalized collision energy 26. Dynamic exclusion was set to 30 s and ions with charge +1, +7, and >+7 were excluded.

MS/MS fragmentation spectra were searched with Proteome Discoverer SEQUEST (version 2.2, Thermo Scientific) against in-silico tryptic digested Uniprot all-reviewed *Homo sapiens* database (release Jun 2017, 42,140 entries) plus all recombinant protein sequences used in this study. The maximum missed cleavages was set to 2. Dynamic modifications were set to oxidation on methionine (M, +15.995 Da) and acetylation on protein N-terminus (+42.011 Da). Carbamidomethylation on cysteine residues (C, +57.021 Da) was set as a fixed modification. The maximum parental mass error was set to 10 ppm, and the MS/MS mass tolerance was set to 0.02 Da. The false discovery threshold was set strictly to 0.01 using the Percolator Node validated by q-value. The relative abundance of parental peptides was calculated by integration of the area under the curve of the MS1 peaks using the Minora LFQ node.

### Data analysis and statistics

The Proteome Discoverer peptide groups abundance values were normalized to the corresponding samples’ median values. After normalization, the missing values were imputed using the K-nearest neighbor algorithm (46). The optimized number of neighbors was determined to be n = 10. The protein copy numbers were assessed using intensity-based absolute quantification (iBAQ) (47). Proteomics data analysis was performed in Microsoft Excel, R (version 3.4.2), and Perseus (version 1.6.2.2).

### Metabolite Set Enrichment Analysis

MCF-10A intracellular pool sizes were ranked based on log_2_ fold change, and enrichment analysis was run with unweighted statistic using the Broad Institute’s GSEA java applet against all KEGG metabolic pathways. Statistical significance was assessed by 5,000 permutations of the ranked list.

## DATA AVAILABILITY

The mass spectrometry proteomics data have been deposited to the ProteomeXchange Consortium via the PRIDE partner repository (48) with the dataset identifier PXD015122 (reviewer username, reviewer password: X1dvWm5c).

### ACKNOWLEDGMENTS

This work was supported by The Margaret E. Early Medical Research Trust, The Rose Hills Foundation, The USC Provost’s Office, and The Viterbi School of Engineering. D.Z. was supported by a Mork Family Doctoral Fellowship. J.H.S. was supported by the USC Provost’s Undergraduate Research Fellowship. We would like to thank Melanie MacMullan and Dr. Pin Wang for assistance with flow cytometry experiments.

## AUTHOR CONTRIBUTIONS

D.Z. and N.A.G. conceived the project. D.Z., J.H.S., M.P.J., and S.T.P. conducted the experiments. A.D. developed the LC-MS metabolomics methodology. D.Z., J.H.S., M.P.J., and N.A.G. interpreted data, and D.Z. and N.A.G. wrote the manuscript.

