## Supplemental Figures for "AKT but not MYC promotes reactive oxygen species-mediated cell death in oxidative culture"

Running title: AKT promotes oxidative stress in galactose culture

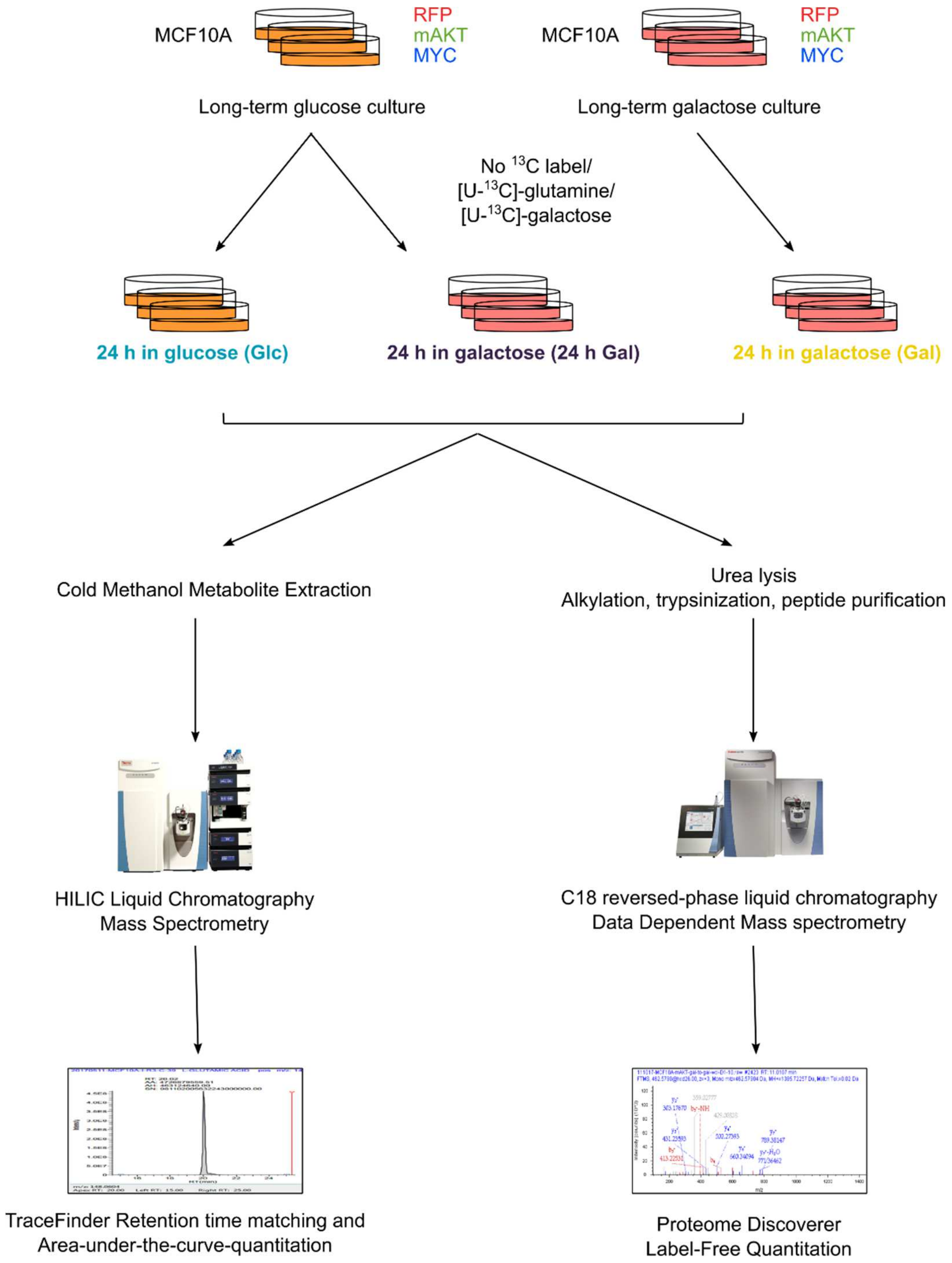


**Supp. Figure S1. Experimental overview**

MCF-10A cells expressing either RFP, mAKT, or MYC were cultured in glucose before switching to glucose (Glc) or galactose media for 24 h (24 h Gal). Long-term galactose cultured MCF-10A RFP, AKT and MYC cells (>5 passages in galactose) were cultured in galactose before switching to galactose (Gal) containing media for 24 h. For metabolomics, intracellular metabolites were extracted using cold methanol and then analyzed using hydrophilic interaction liquid chromatography (HILIC)-mass spectrometry (1,2). For proteomics, cells were lysed in 8M urea, followed by reduction, alkylation, and trypsinization. We performed two independent biological replicates, and each experiment were subjected to two technical LC-MS injections. Protein levels were calculated using label-free quantitation and the iBAQ method (3).

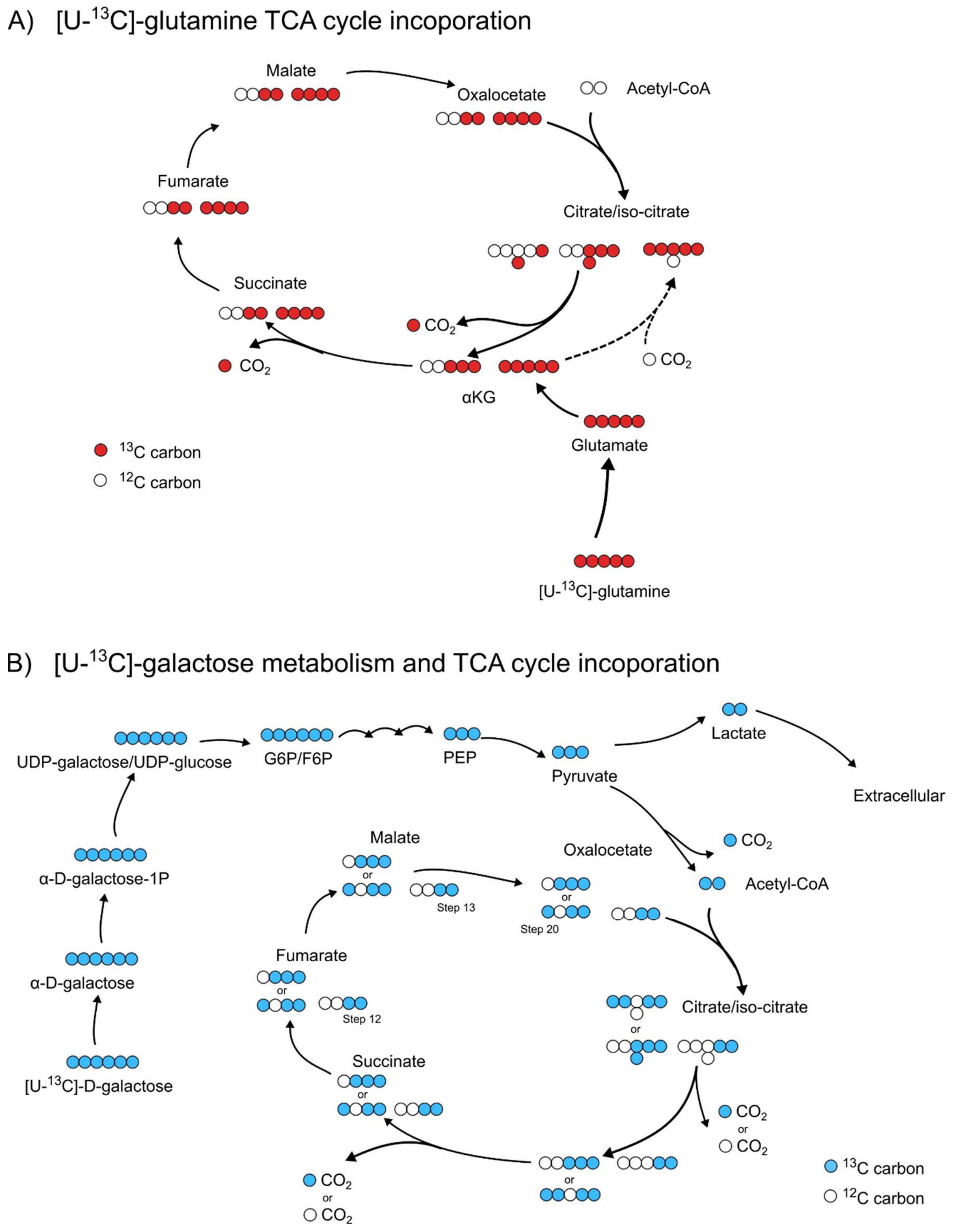


**Supp. Figure S2. Atom transition maps for [U-$^{13}$C]-L-glutamine and [U-$^{13}$C]-galactose stable isotope labeling.**

**A)** [U-$^{13}$C]-L-glutamine is converted to glutamate and then α-ketoglutarate (αKG) to enter the TCA cycle. Solid arrows indicate forward TCA cycle activity and dashed arrows indicate reductive carboxylation (4).

**B)** [U-$^{13}$C]-D-galactose enters cellular metabolism by isomerization to α-D-galactose, conversion to galactose-1-phosphate (α-D-galactose-1P) and UDP-galactose/UDP-glucose before conversion to glucose-6-phosphate (G6P). G6P can then enter glycolysis and the TCA cycle (5).

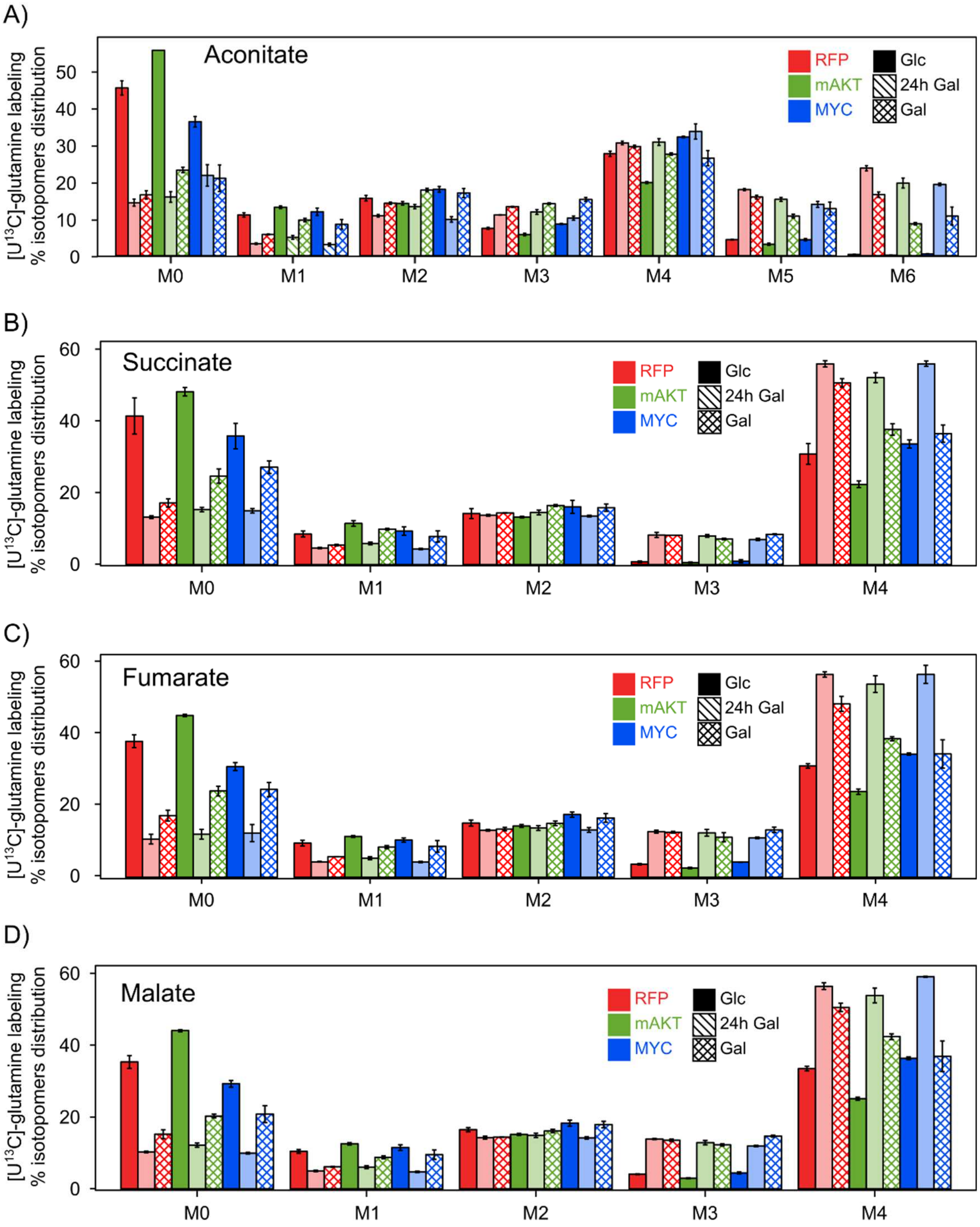


**Supp. Figure S3. Differential usage of glutamine in mAKT-expressing cells.**

$^{13}C$ isotopomer distributions from MCF-10A cells labeled with [U-$^{13}C$]-L-glutamine. Isotopomer abundances were normalized to the sum of all isotopomers to calculate the percent abundance of each isotopomer. **A)** aconitate; **B)** succinate; **C)** fumarate; and **D)** malate from the TCA cycle show that AKT cells in glucose condition were more dependent on non-glutamine sources whereas MYC cells were more dependent on glutamine-derived carbon. This suggests that MYC cells should have less problem switching to galactose. Increasing incorporation of M5 citrate-isocitrate suggest all three cells type were forced to utilize $^{13}C$-glutamine to produce citrate-isocitrate through reductive carboxylation

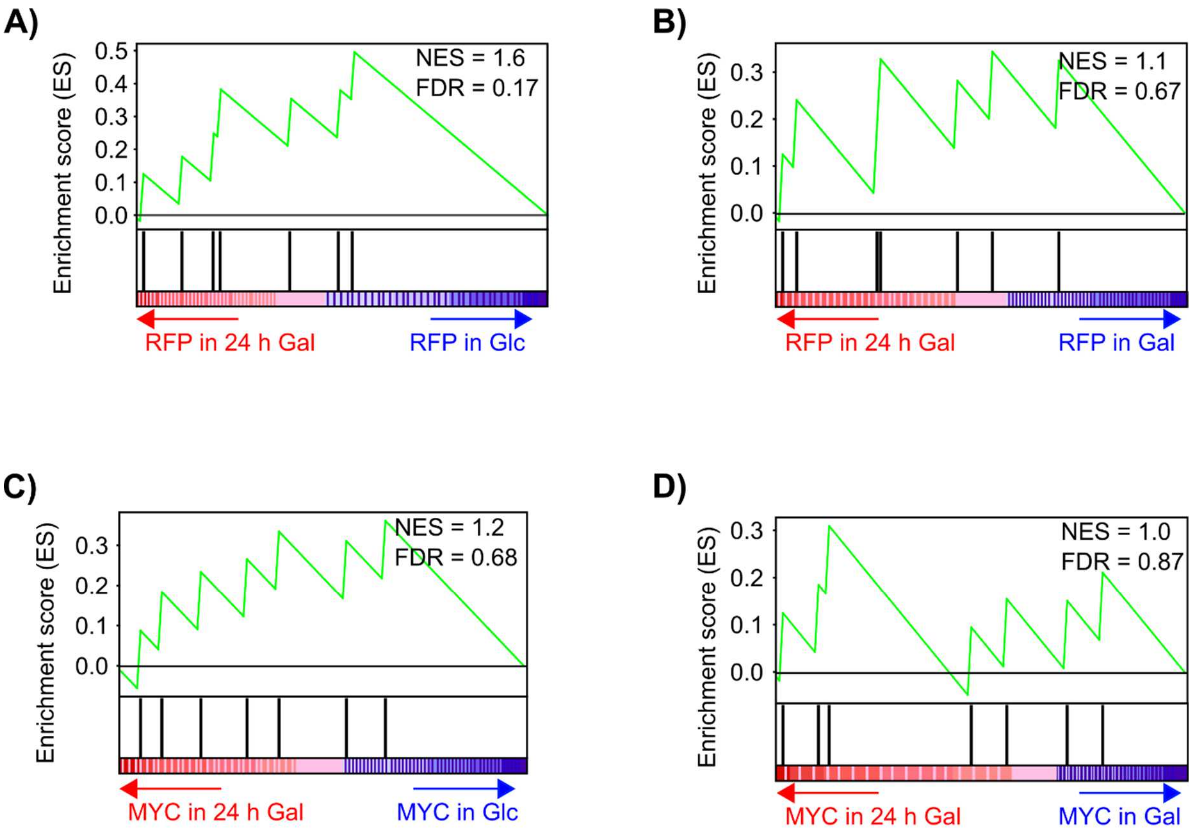


**Supp. Figure S4. Metabolite set enrichment analysis of glutathione metabolism for RFP- and MYC-expressing MCF-10A cells.**

Metabolite set enrichment analysis mountain plots for glutathione metabolism for **A)** RFP-expressing MCF-10A cells comparing short-term galactose culture to glucose culture (24 h Gal/Glc), **B)** RFP-expressing cells comparing short-term to long-term galactose culture (24 h Gal/Gal), **C)** MYC-expressing MCF-10A cells comparing short-term galactose culture to glucose culture (24 h Gal/Glc), and **D)** MYC-expressing cells comparing short-term to long-term galactose culture (24 h Gal/Gal). The green line denotes the enrichment score, and the black tick marks denote metabolites that belong to glutathione metabolism. The normalized enrichment score (NES) and false discovery rate (FDR) are shown. In comparison to mAKT-expressing cells (Fig. 3), RFP- and MYC-expressing cells did not exhibit upregulation of the glutathione metabolism pathway.

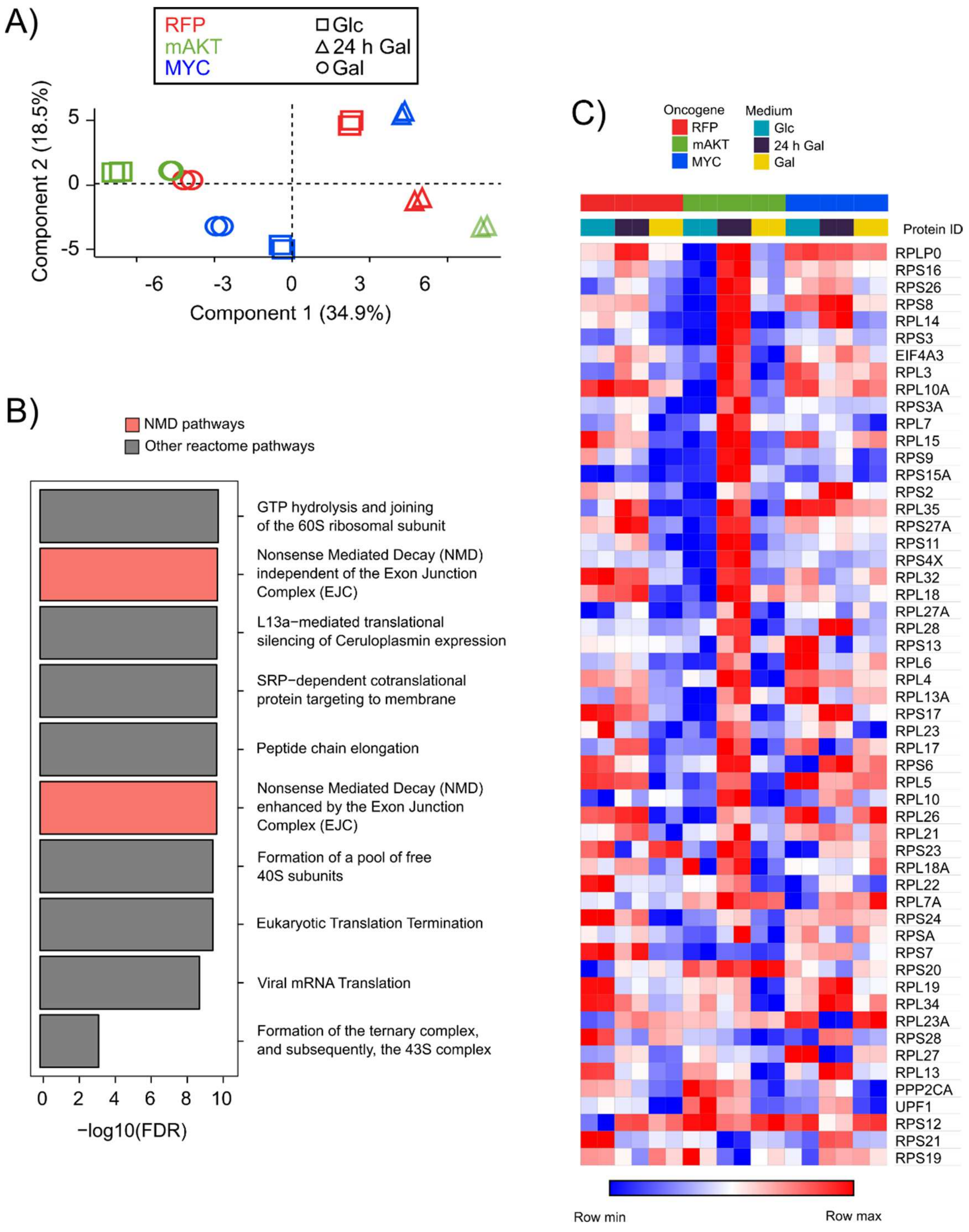


**Supp. Figure S5. LC-MS proteomics Experiment 2 demonstrates enrichment of nonsense-mediated mRNA decay (NMD) proteins in mAKT cells switched to galactose.**

**A)** Principal component analysis score plots (PC1 vs. PC2) of proteomic data from Experiment 2 segregated samples by oncogene and media type. Color denotes oncogene, and shape denotes media type. Each sample was analyzed in technical duplicate. Short-term galactose culture (24 h Gal) induced a positive shift on PC1 (34.9% of variation) for all cell types relative to glucose culture (Glc). Long-term galactose culture (Gal), in contrast, exhibited a negative shift on PC1 relative to short-term galactose culture. The PC1 shift for mAKT-expressing cells was significantly larger than for either RFP- or MYC-expressing cells. Similar trends were seen in Experiment 1 (Fig. 4B).

**B**) Enrichment analysis identified Reactome pathways enriched in the PC1 loadings vector from Experiment 2. Nonsense Mediated Decay (NMD) enhanced by the Exon Junction Complex (EJC) and Nonsense Mediated Decay (NMD) independent of the Exon Junction Complex (EJC) are highlighted. Similar trends were seen in Experiment 1 (Fig. 4C)

**D**) mAKT-expressing cells significantly upregulated NMD proteins in short-term galactose culture. A heatmap of protein expression from Experiment 2 for the Nonsense Mediated Decay (NMD) enhanced by the Exon Junction Complex (EJC) pathway (Reactome R-HSA-975957) demonstrated that mAKT-expressing cells dramatically upregulated NMD protein expression in short-term (24 h) galactose culture. Similar trends were seen in Experiment 1 (Fig. 4D).

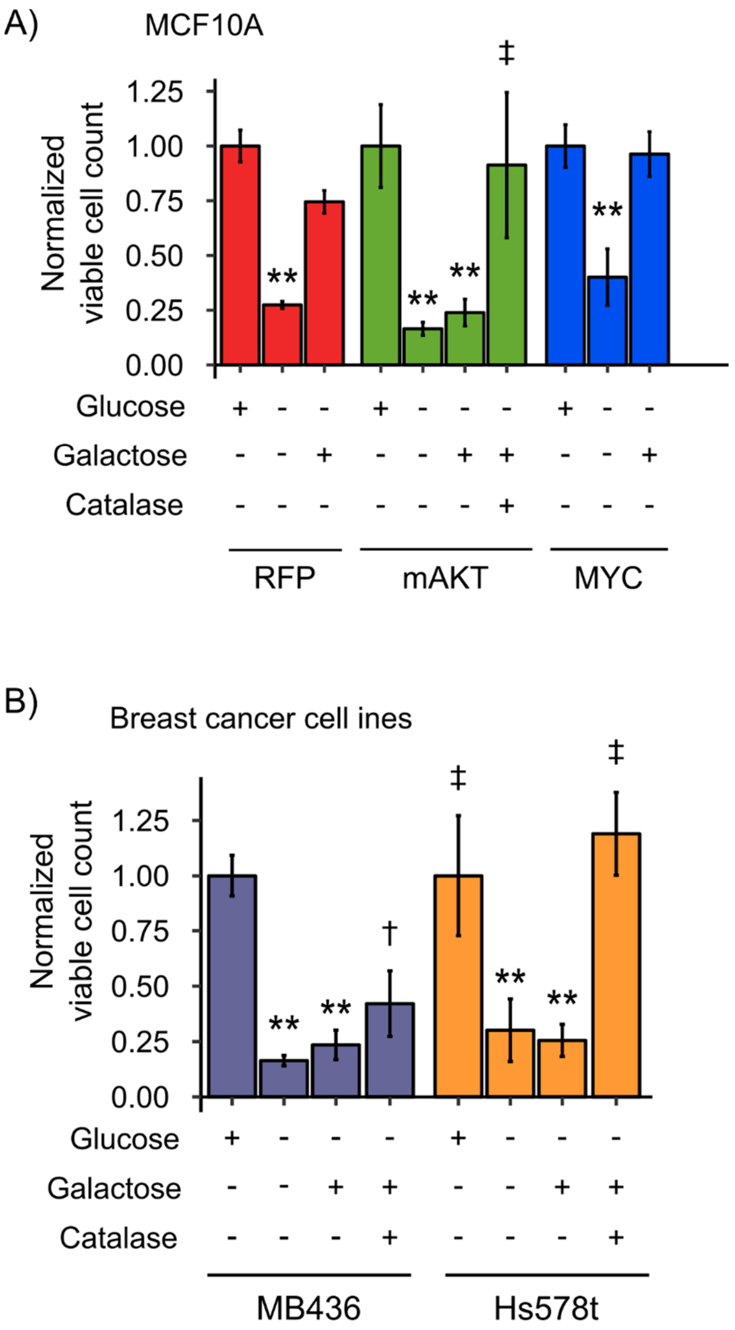


**Supp. Figure S6. ROS are required for galactose-induced cell death.**

**A**) The ROS scavenger catalase rescued mAKT-expressing cells from galactose-induced cell death. MCF-10A cells expressing RFP, mAKT, or MYC were cultured in glucose, without glucose, or in galactose with or without 200 U/ml of the ROS scavenger catalase for 32 h. Viable cell number was measured by trypan blue staining and normalized to the glucose cultured cells. ** denotes Student's t-test p-value < 0.01 compared to glucose culture, and ‡ denotes Student's t-test p-value < 0.01 compared to galactose culture without catalase (n=2-4 biological replicates).

**B**) The ROS scavenger catalase rescued MB436 and Hs578t breast cancer cells from galactose-induced cell death. MB436 and Hs578t cells were cultured in glucose, without glucose, or in galactose with or without 200 U/ml of the ROS scavenger catalase for 24 h. Viable cell number was measured by trypan blue staining and normalized to the glucose cultured cells. ** denotes Student's t-test p-value < 0.01 compared to glucose culture, and ‡ denotes Student's t-test p-value < 0.01 compared to galactose culture without catalase (n=3-4 biological replicates).
